# Subject classification and cross-time prediction based on functional connectivity and white matter microstructure features in a rat model of Alzheimer’s using machine learning

**DOI:** 10.1101/2023.03.27.534331

**Authors:** Yujian Diao, Ileana Ozana Jelescu

## Abstract

**Background:** The pathological process of Alzheimer’s disease (AD) typically takes up decades from onset to clinical symptoms. Early brain changes in AD include MRI-measurable features such as aItered functional connectivity (FC) and white matter degeneration. The ability of these features to discriminate between subjects without a diagnosis, or their prognostic value, is however not established.

**Methods:** The main trigger mechanism of AD is still debated, although impaired brain glucose metabolism is taking an increasingly central role. Here we used a rat model of sporadic AD, based on impaired brain glucose metabolism induced by an intracerebroventricular injection of streptozotocin (STZ). We characterized alterations in FC and white matter microstructure longitudinally using functional and diffusion MRI. Those MRI-derived measures were used to classify STZ from control rats using machine learning, and the importance of each individual measure was quantified using explainable artificial intelligence methods.

**Results:** Overall, combining all the FC and white matter metrics in an ensemble way was the best strategy to discriminate STZ rats, with a consistent accuracy over 0.85. However, the best accuracy early on was achieved using white matter microstructure features, and later on using FC. This suggests that consistent damage in white matter in the STZ group might precede FC. For cross-timepoint prediction, microstructure features also had the highest performance while, in contrast, that of FC was reduced by its dynamic pattern which shifted from early hyperconnectivity to late hypoconnectivity.

**Conclusions:** Our study highlights the MRI-derived measures that best discriminate STZ vs control rats early in the course of the disease, with potential translation to humans.

## Background

Alzheimer’s disease (AD) as a progressive neurodegenerative disorder is the main cause of dementia, which is characterized by decline in cognitive function such as thinking, remembering, and reasoning. AD can be divided into two major categories: sporadic AD and familial AD. The familial AD that accounts for less than 5% of all AD cases (Bali et al., 2012) is usually caused by a genetic mutation, whereas sporadic AD accounting for the majority of AD cases is multifactorial (Castanho and Lunnon, 2019). Pathologically, AD is characterized by extracellular deposits of Aβ peptides as senile plaques, intraneuronal neurofibrillary tangles, reduced brain glucose metabolism and large-scale neuronal loss in the most affected regions of the brain, such as the medial temporal lobe and neocortical structures (Breijyeh and Karaman, 2020; De-Paula et al., 2012; Du et al., 2018; Liu et al., 2019; Long and Holtzman, 2019).

Non-invasive brain imaging techniques such as magnetic resonance imaging (MRI) play a vital role in detecting early changes in the brain associated with AD. Gross cerebral atrophy (Gispert et al., 2015), white matter (WM) degeneration (Agosta et al., 2011; Chang et al., 2015; Choo et al., 2010; Dong et al., 2020; Jelescu et al., 2018) and altered functional connectivity (FC) (Agosta et al., 2012; Binnewijzend et al., 2012; Brier et al., 2012; Franzmeier et al., 2019; Greicius et al., 2004) were found to be relevant biomarkers. Recently, resting-state FC has been proposed to identify individuals at risk for Alzheimer’s disease in the early stages (Hojjati et al., 2019; Zamani et al., 2022; Zhang et al., 2019). The characterization of the temporal progression of microstructural and FC changes promises to provide an understanding of disease mechanisms, an effective disease staging and a window for therapeutic intervention.

As the pathological cascade of AD takes up to years or even decades from the dementia onset to full-blown manifestations, it remains challenging to acquire comprehensive longitudinal data on prospective AD subjects. As an alternative, animal models can be valuable tools to obtain data across the lifespan and study each of the contributors to the AD cascade individually, thus untangling direct effects of contributors and their interactions. Although numerous animal models have been developed to replicate the AD phenotype, most of them are transgenic models which are less representative of sporadic AD and are primarily based on the Aβ hypothesis (Buxbaum, 2009; King, 2018), which is increasingly challenged (Kametani and Hasegawa, 2018). However, with glucose hypometabolism being increasingly recognized as a potential cause of AD (Correia et al., 2011; Hölscher, 2019; Kuehn, 2020), animal models of brain insulin resistance have been developed by an intracerebroventricular (icv) injection of streptozotocin (STZ) (Knezovic et al., 2015; Kraska et al., 2012; Shoham et al., 2003). The icv-STZ animals have been reported to manifest typical features of AD such as memory impairment, extracellular accumulation of Aβ, neuronal loss, axonal damage and demyelination in the hippocampus and fimbria (Shoham et al., 2003), reduced glucose uptake (Heo et al., 2011), and oxidative stress (N. Lester-Coll et al., 2006; Shoham et al., 2003), without developing systemic diabetes.

In a previous work, we performed a comprehensive longitudinal study (Tristão Pereira et al., 2021) in a icv-STZ rat model to quantitatively characterize alterations in FC and in WM microstructure using resting-state functional MRI (fMRI) and advanced diffusion MRI techniques, respectively, as well as in brain glucose uptake captured by ^18^FDG-PET. By comparing the STZ group to the control group, non-invasive MRI-derived measures of functional breakdown and WM degeneration were identified and evaluated in the context of brain glucose hypometabolism. Alterations in resting-state FC in STZ rats were found in brain regions closely associated with AD (Agosta et al., 2012; Brier et al., 2012) with broadly increased then decreased connectivity at early and late timepoints, respectively. WM microstructure metrics derived from DKI (an extension of Diffusion Tensor Imaging – DTI – that provides complementary information about tissue heterogeneity (Jensen et al., 2005)) and the WMTI-Watson biophysical model (Fieremans et al., 2011; Jespersen et al., 2018) revealed specifically intra-axonal damage and axonal loss in the corpus callosum, fimbria and cingulum of STZ rats. The temporal dynamics of both WM integrity and FC were consistent with previously reported nonmonotonic trajectories of brain alterations along AD progression in humans (Dickerson et al., 2005; Dong et al., 2020; Pegueroles et al., 2017; Schultz et al., 2017). These findings not only reinforced the suitability of the icv-STZ animal model for sporadic AD but also proposed MRI-derived features to identify alterations in the prodromal stage and monitor disease progression.

In this study, we go beyond descriptive statistics and evaluate the microstructural and functional measures for their potential to discriminate between control and STZ groups at a given timepoint and across time. We utilize them as features using machine learning (ML) to train classification models such as Logistic Regression (LR) to differentiate individual subjects. Moreover, we employ explainable artificial intelligence methods to interpret ML model outcomes. For example, the importance of each feature in terms of the absolute value of LR coefficients is used to identify features best discriminating STZ rats from controls. SHAP values (SHapley Additive exPlanations) (Lundberg and Lee, 2017), a model-agnostic approach, are used to interpret the model outcomes and to improve model transparency. Finally, the dynamic relationships between the functional and microstructural measures in STZ rats are highlighted at the early and late timepoints of disease progression. In a nutshell, our study highlights the MRI-derived measures that best discriminate STZ vs control rats at various stages of the disease, with potential translation to humans.

## Methods

### Study design

Male Wistar rats (236 ± 11 g) underwent a bilateral icv-injection of either streptozotocin (3 mg/kg, STZ group) or buffer (CTL group) as previously described (Tristão Pereira et al., 2021). When delivered exclusively to the brain, streptozotocin induces impaired brain glucose metabolism and is used as a model of sporadic AD (Grieb, 2016; Nataniel Lester-Coll et al., 2006). Resting-state fMRI and diffusion MRI data were acquired longitudinally at four timepoints (2, 6, 13 and 21 weeks since icv injection) (**Figure 1**). Timepoints were chosen to be consistent with previous rat STZ studies while also accommodating for constraints of repeated MRI scanning related to anesthesia, cannulations, and scanner availability.

**Figure 1.**
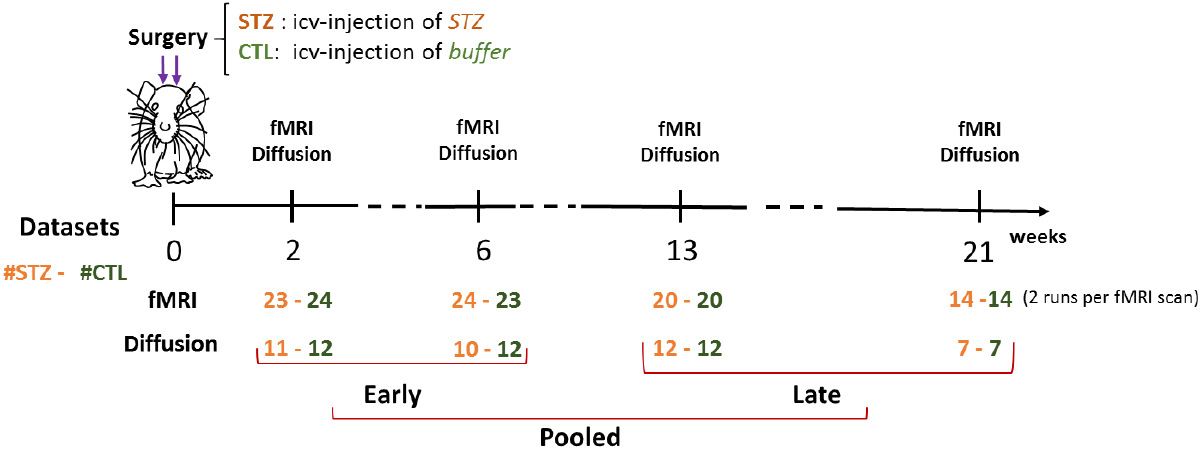
Experimental timeline. FMRI and diffusion MRI data were collected at 2, 6, 13 and 21 weeks after icv-STZ injection. Two fMRI runs were acquired for each MRI session. The 4 timepoints were further grouped into *Early* and *Late* time groups and finally the *Pooled* dataset. Sample sizes (STZ/CTL) in the 3 datasets for fMRI and diffusion MRI are as follows. Early: 47/47 (fMRI), 21/24 (diffusion); Late: 34/34 (fMRI), 19/19 (diffusion); Pooled: 81/81 (fMRI), 40/43 (diffusion).

### MRI data acquisition

Animals were initially anesthetized using isoflurane (4% for induction and 1–2% for maintenance in an oxygen/air mixture of 30%/70%) and positioned in a homemade MRI cradle equipped with a fixation system (bite bar and ear bars). A catheter was inserted subcutaneously on the back of the animal for later medetomidine delivery. One hour before starting the resting-state fMRI acquisition, anesthesia was switched from isoflurane to medetomidine (Dorbene, Graeub, Switzerland), which preserves neural activity and vascular response better than isoflurane (Pawela et al., 2009; Weber et al., 2006), with an initial bolus of 0.1 mg/kg followed by a continuous perfusion of 0.1 mg/kg/h (Reynaud et al., 2019). The commercial solution at 1 mg/mL was diluted to 0.033 mg/mL. Throughout the experiment, the breathing rate was monitored using a respiration pillow and a rectal thermometer, respectively. Body temperature was maintained around (37±0.5) °C. The breathing rate under medetomidine was around 85 bpm. At the end of the scanning session, animals were woken up with an intramuscular injection of atipamezole (Alzane, Graeub, Switzerland) at 0.5 mg/kg.

MRI experiments were conducted on a 14.1 T small animal scanner. As a result of system upgrade, data were acquired with two different consoles for the magnet: Varian system (Varian Inc.) equipped with 400 mT/m gradients (Cohort 1, N=17 rats) and Bruker system (Ettlingen, Germany) equipped with 1 T/m gradients (Cohort 2, N=7 rats), both using the same in-house built quadrature surface transceiver. The acquisition parameters were the same for the two cohorts. Each cohort comprised animals from both groups: Cohort 1 (CTL/STZ, N = 8/9 rats) and Cohort 2 (CTL/STZ, N = 4/3 rats).

Structural *T*_2_-weighted images were collected using a fast spin-echo sequence with the following parameters: TE/TR = 10.17/3000 ms, echo train length: 4, matrix size = 128 × 128, FOV = 19.2 × 19.2 mm^2^, voxel size = 0.15 × 0.15 mm^2^, 30 coronal 0.5-mm slices, scan time = 10 minutes.

Diffusion-weighted data were acquired using a pulsed-gradient spin-echo segmented echo-planar-imaging (EPI) sequence, with the following protocol: 4 b = 0 images and 3 shells at b = 0.8 / 1.3 / 2.0 ms/µm^2^, with 12, 16 and 30 directions, respectively; δ/Δ = 4/27 ms; TE/TR = 48/2500 ms; 4 shots; matrix size = 128 × 64, Field-of-view = 23 × 17 mm^2^, voxel size = 0.18 × 0.27 mm^2^, 9 coronal 1-mm slices, 4 repetitions, scan time = 1 hour. Resting-state fMRI data were acquired using a two-shot gradient-echo EPI sequence as follows: TE/TR = 10/800 ms, TRvolume = 1.6 s, matrix size = 64 × 64, Field-of-view = 23 × 23 mm^2^, voxel size = 0.36 × 0.36 mm^2^ and 8 coronal 1.12-mm slices, 370 repetitions, scan time = 10 minutes. Two fMRI runs were acquired in each MRI session.

### Data processing

FMRI data processing followed the PIRACY pipeline (Diao et al., 2021) which included denoising (Veraart et al., 2016), susceptibility distortion correction (Smith et al., 2004), slice-timing correction (Henson et al., 1999), spatial smoothing, and removal of physiological noise following independent component (IC) analysis decomposition. FC matrices between 28 regions of interest (ROIs) based on the Waxholm Space Atlas were computed, co-varying for the global signal (Diao et al., 2021). Statistical comparisons of FC between the STZ and CTL groups at each timepoint were performed using NBS (Zalesky et al., 2010) to identify network connections that showed significant between-group differences. Specifically, NBS uses one-tailed two-sample *t*-test to detect differences in group averaged FC between the two groups. Thereby, two contrasts (STZ > CTL and STZ < CTL) were tested separately. A *t*-statistic threshold of 2.2 was chosen on the basis of medium-to-large sizes of the subnetwork comprised connections with their *t*-statistic above the threshold (Tsurugizawa et al., 2019) as well as the underlying *p*-values. Significance (*p* ≤ 0.05) was tested after family wise error rate correction using non-parametric permutation (N = 5000).

Diffusion data processing included MP-PCA denoising (Veraart et al., 2016), Gibbs-ringing correction (Kellner et al., 2016) and correction for susceptibility distortions and eddy currents using FSL’s eddy (Andersson and Sotiropoulos, 2016). The diffusion and kurtosis tensors were estimated using a weighted linear least squares algorithm (Veraart et al., 2013) and typical DKI-derived metrics were computed: fractional anisotropy (FA), mean/axial/radial diffusivity and mean/axial/radial kurtosis. The biophysical WMTI-Watson model (Jespersen et al., 2018) (**Figure 2**) was estimated voxel-wise in WM regions using nonlinear least squares fitting to extract its microstructure parameters: *f, D*_*a*_, *D*_*e*,∥_, *D*_*e*,⊥_, and *c*_2_. In parallel, fractional anisotropy (FA) maps were registered to an FA template in the Waxholm Space using linear and non-linear registration in FSL (Jenkinson et al., 2002) and the corpus callosum (CC), cingulum (CG) and fimbria (Fi) of the hippocampus were automatically segmented. For each ROI, average tensor and biophysical model metrics were calculated. Group differences were tested using *t*-test.

**Figure 2.**
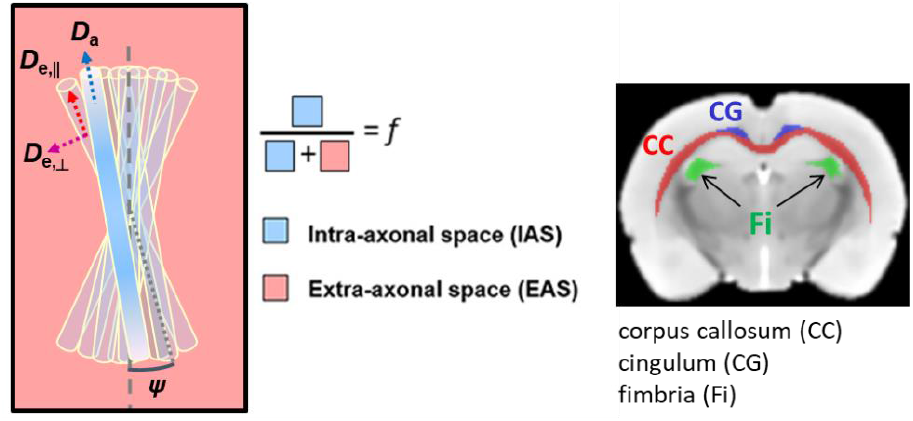
Schematic of the WMTI-Watson biophysical model and the white matter ROIs. The diffusion signal is described in terms of two non-exchanging compartments, the intra and extra-axonal spaces. Here, the axons are modelled as sticks with radius equal to zero. The intra-axonal space is described by a relative volume fraction of water *f* and by the parallel intra-axonal diffusivity *D*_*a*_. The perpendicular intra-axonal diffusivity is negligible at the relevant diffusion times and weightings. The bundle of axons is embedded in the extra-axonal space, characterized by its parallel *D*_*e*,∥_ and perpendicular extra-axonal diffusivities *D*_*e*,⊥_. The axons’ orientations are modeled by a Watson distribution, which is characterized by⟨ (*cosψ*)^2^ ⟩≡ *c*_2_.

### Classification using logistic regression

For FC-based classification, correlation coefficients between ROIs in the FC matrix were taken as classification features by vectorizing the upper triangle of the FC matrix since FC is symmetric. To study the connection between statistical differences and classification performance in discriminating the two groups, significant edges from the NBS analysis were selected as a reduced list of features for classification. Datasets were grouped as Early (2 & 6 weeks, N=94) and Late timepoint (13 & 21 weeks, N=68), as well as all timepoints (Pooled, N=162). STZ/CTL classification using a LR model was trained and tested on each subset (Pooled, Early, and Late), which was normalized to [-1, 1] and randomly split into training (70%) and test datasets (30%). Since the data size was relatively small, the procedure of data splitting, training and testing was repeated 1’000 times and results were aggregated in order to reduce bias.

For microstructure-based classification, there were two types of features for each of the three WM ROIs: I) DKI tensor metrics including FA, axial, mean and radial diffusivities (AxD, MD, RD), axial, mean and radial kurtosis (AK, MK, RK); II) WMTI-Watson model parameters including *f, D*_*a*_, *D*_*e*,∥_, *D*_*e*,⊥_ and *c*_2_ (**Figure 2**). These two kinds of features were used in two ways: as independent feature sets (i.e., DKI only and WMTI only) and combined as a single feature set. As for FC, diffusion datasets were grouped as Early (2&6 weeks, *N*=45), Late (13&21 weeks, *N*=38) and all timepoints (Pooled, *N*=83). LR models of STZ/CTL classification were trained and tested on the three datasets independently with 70% data for training and 30% for testing. The procedure was also repeated 1’000 times.

Considering the small data size, feature dimensionality reduction was also tried for each classification by employing the principal component analysis (PCA) with various number of components.

Morever, we tested classifying STZ and CTL rats by combining the FC and WM microstructure metrics in two distinct ways. One was to create a single classifier based on the concatenation of features of FC and microstructure metrics. The second way was using ensemble learning (Opitz and Maclin, 1999; Rokach, 2010) where three independent classifiers were built each based on one of the three types of features (FC, DKI and WMTI). Their predictions for each class were aggregated and the class with the majority vote was retained. Datasets for which both FC and dMRI were not available jointly (e.g. as a result of partial or artefacted data) were removed, resulting in a slightly reduced sample size (STZ/CTL=38/41).

Finally, cross prediction was performed, which means a classifier was trained on the dataset of one timepoint (e.g. Early) and tested on the other timepoint (e.g. Late) and vice versa. Cross prediction was tested on classifiers built on both separate and joint features.

### Model explainability and feature importance

Classification accuracy was used to assess the performance of a LR model in classifying STZ and CTL rats. However, to better interpret and explain the model outcome, we further calculated the importance of each feature in driving a model to predict the STZ class in terms of the absolute values of LR coefficients (Smerdov et al., 2019). The mean feature importance was computed by averaging the absolute LR coefficient of each feature over the 1’000 repetitions of training/test data splits, along with mean classification accuracy and standard deviation.

SHAP values that have been widely used for interpreting ML models were also calculated. It is a game theory-based approach used to measure the contribution of each player in a cooperative game by aggregating marginal contributions brought by a feature to various kinds of feature combinations. In this study, SHAP values were computed for different types of features (i.e. significant FC connections, DKI metrics or WMTI parameters) at each of the three timepoints (early, late and pooled) to measure the individual impact of each feature on the model outcome. SHAP values indicated which features drove a model to predict the STZ or CTL class. With the combination of classification accuracy and SHAP values, we were able to validate each measure’s ability to discriminate STZ rats from controls. As SHAP values are instance-based, they cannot be averaged over repeated training scenarios like the LR coefficient. Instead, we selected representative SHAP value sets from the 1’000 candidates by choosing the ones that had relatively high classification accuracies (> 0.9) in both training and test datasets such that the model would have good performance as well as high generalizability.

## Results

### FC-based classification

In **Figure 3**, graph networks highlight the group differences in nodal connections in the Pooled, Early, and Late datasets. Up to six weeks after icv injection (Early), STZ group displayed increased connectivity within the default mode network (DMN) (including the the anterior cingulate cortex (ACC), retrosplenial cortex (RSC), hippocampus and subiculum) as well as striatum and decreased connectivity between the DMN (RSC, posterior parietal cortex – PPC – and hippocampus) and the lateral cortical network including primary and secondary somatosentory cortex (S1,S2) and the motor cortex, as compared to CTL rats. From 13 weeks on (Late), reduced connectivity became more widespread within the DMN and lateral cortical network in STZ rats.

**Figure 3.**
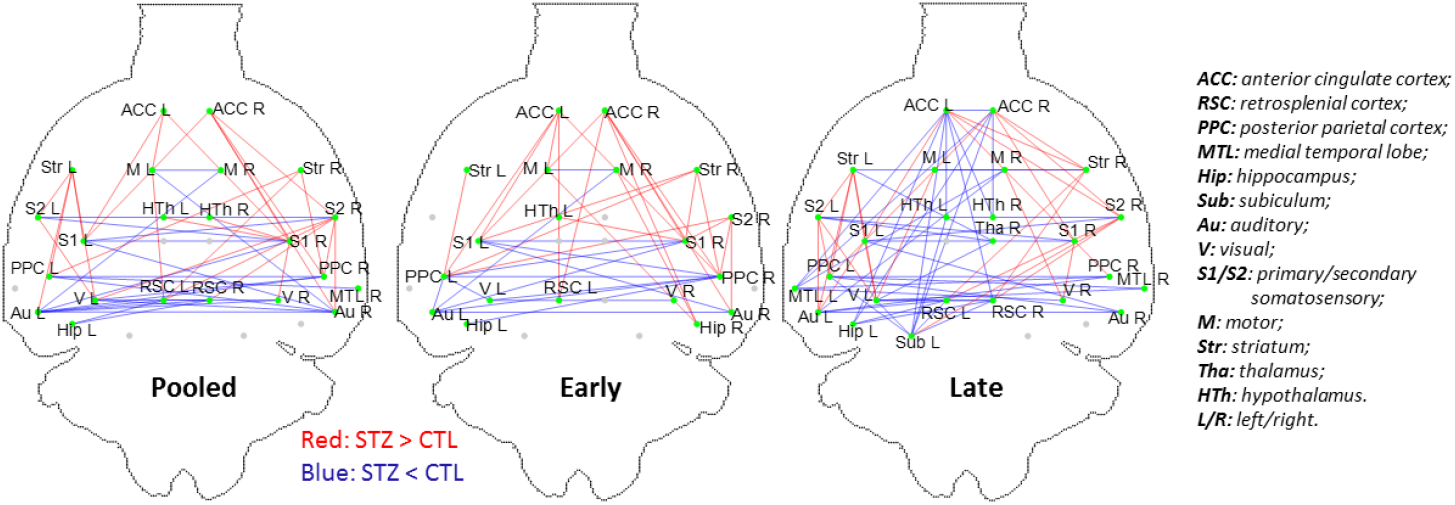
Graph networks of significant group difference using NBS with *p* < 0.05 (family-wise error rate corrected) for the 3 datasets (Pooled, Early and Late). Blue/red edges represent edges where STZ rats have weaker/stronger FC than CTL.

When using all connections as features (N=378), prediction accuracy on the Pooled, Early, and Late datasets was 0.75, 0.69 and 0.83, respectively. The most relevant edges involved the ACC, hypothalamus, RSC, hippocampus and subiculum as nodes (**Figure 4**A), in agreement with edges found as significantly different between groups in the NBS analysis (**Figure 3**). When only significant edges from the NBS analysis were selected as a reduced list of features for classification (N=49, 38 and 71 features in the Pooled, Early, and Late datasets, respectively), the classification accuracy improved to 0.79 for Pooled, 0.72 for Early, and 0.90 for Late datasets (**Figure 4**B). Improved accuracy was not strictly related to feature reduction: reducing features using PCA deteriorated classification accuracy (data not shown). Notably, the highest classification accuracy was found on the Late dataset which is consistent with the advanced stage of disease and more marked differences between STZ and CTL.

**Figure 4.**
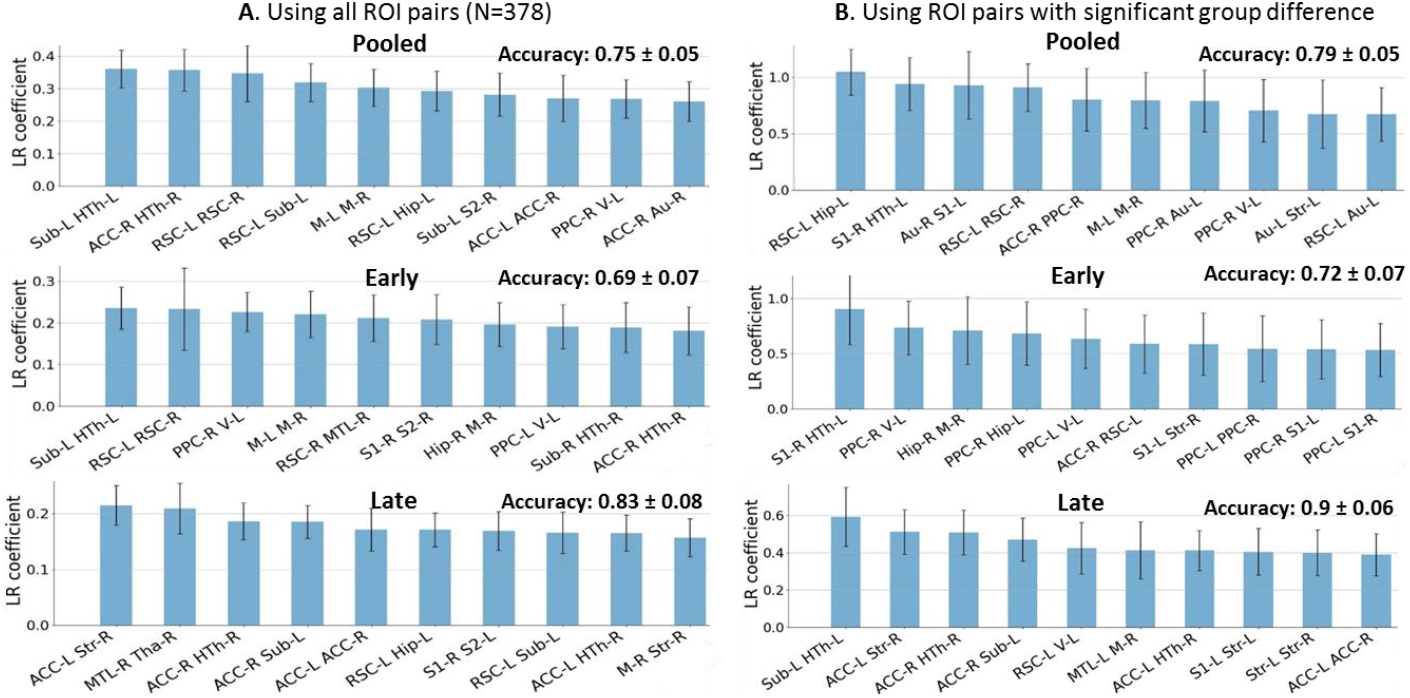
A) Top ten features (out of 378) and their importance in terms of absolute LR coefficient in rat classification on the FC dataset (mean ± standard deviation, averaged over 1000 repetitions). Each feature is an edge. The most relevant edges that discriminate between CTL and STZ rats involve ACC, hypothalamus (HTh), RSC, hippocampus (Hip) and subiculum (Sub). B) Only connections surviving NBS significance test were selected as features for classification (top 10 displayed). Classification accuracy was improved from 0.75 to 0.79 for Pooled, 0.69 to 0.72 for Early and 0.83 to 0.90 for Late dataset by this feature pre-selection. Higher classification accuracy in Late dataset is consistent with the advanced stage of disease and more marked differences between STZ and CTL.

However, the top 10 edges with the highest feature importance in the first classification (all features, **Figure 4**A) did not overlap strongly with that from the second classification (reduced features, **Figure 4**B) perhaps due to the small sample size. The nodes involved in the top 10 edges did however overlap strongly between the two classifications.

**Figure 5** displays SHAP plots for each instance of the most important features used to classify STZ and CTL subjects in each of the three datasets. Top features were generally consistent with those from LR in **Figure 4**A. Distribution of values for each feature (edge) in the STZ and CTL groups also agreed with the group difference test in the form of graph networks (**Figure 3**). For example, in the early timepoints, both methods revealed the STZ group had stronger connectivity between right hippocampus and motor cortex, left ACC and S1, left hypothalamus and right S1, and reduced connectivity between right PPC and left visual cortex, as well as right PPC and left hippocampus. In the late timepoints, the STZ group had increased connectivity between left S2 and striatum, left ACC and right striatum, left S1 and striatum, and weaker connectivity between left subiculum and hypothalamus, left RSC and visual cortex. Overall, the distribution of SHAP values demonstrated that the STZ group had hyperconnectivity in the early timepoint but hypoconnectivity in the late timepoint, which confirmed the finding in the previous study (Tristão Pereira et al., 2021).

**Figure 5.**
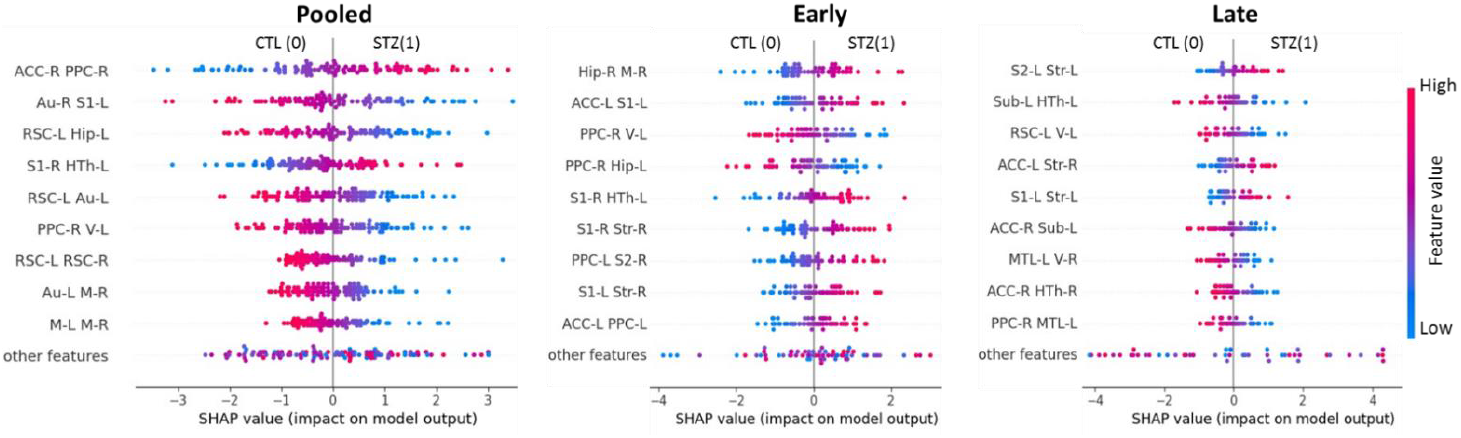
Exemplary SHAP summary plots for the three datasets (Pooled, Early and Late) based on the model using FC significant connections as features. The summary plot combines feature importance with feature effects. Each point on the summary plot is a SHAP value for a feature and an instance. The position on the y-axis is determined by the feature and on the x-axis by the SHAP value. The color represents the value of the feature from low (blue) to high (red). The features are ordered according to their importance (top 9 displayed). Positive SHAP values lead the model to predict 1 (STZ) while negative ones lead the model to predict 0 (CTL).

### Microstructure-based classification

For classification based on WM microstructure features, the mean test accuracy and top features with the highest importance are displayed in **Figure 6**. When using the combined diffusion metrics (DKI+WMTI) as features, the FA in fimbria and corpus callosum stood out as the best discriminating features in the early timepoints while the axonal density (*f*) of the WMTI-Watson model in the fimbria was the most important feature in the late timepoints as well as in the pooled data. Overall, fimbria microstructure was the best discriminator between groups. FA was sensitive to early changes in STZ rats, which drove the DKI-based model to achieve better classification accuracy than the WMTI-based model at the Early timepoint (**Table 1**). While the accuracy of the DKI-based classifier decreased significantly at the Late timepoint, the accuracy of the WMTI-based classifier remained stable across time. The classifier built on combined DKI+WMTI metrics obtained the highest accuracy in the Early stage and similar accuracy in the Late.

**Figure 6.**
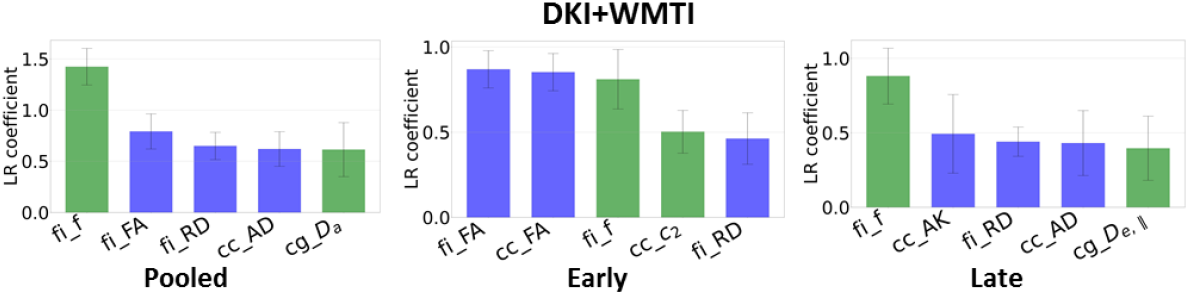
Feature importance and test classification accuracy using different microstructure metrics (mean ± std over 1000 repetitions). Displayed are top 5 most import features on the three datasets using DKI metrics (blue) and WMTI parameters (green) altogether. *fi = fimbria, cc = corpus callosum, cg = cingulum, FA = fractional anisotropy, AD/RD = axial/radial diffusivity, AK = axial kurtosis, f = axonal density, D*_*a*_ *= intra-axonal diffusivity, D*_*e*,||_ *= extra-axonal parallel diffusivity, c*_*2*_: *orientation dispersion*.

**Table 1.** The accuracy of classification on the three datasets and cross predictions in the different cases of employing separate and joint features. The last column is the total data size in the Pooled dataset. The FC dataset has a larger sample size because each rat subject had two fMRI scans for each experiment. Dimension reduction using PCA didn’t improve the classification accuracy in most cases except for FC-based classification on the Early dataset and the late-to-early cross prediction where the new accuracies and the optimal numbers of PCA components were indicated. In FC-based classification, only connections with significant group difference were retained except for cross predictions where all FC connections were used. For Pooled, Early and Late timepoints, the classification accuracy is the average over 1’000 random data splits into 70% training and 30% testing. For Early-to-Late, the training set was all Early datasets and the test set all Late datasets (and vice versa for Late-to-Early).

| Features | Pooled | Early | Late | EarlyToLate | LateToEarly | Sample size |
| --- | --- | --- | --- | --- | --- | --- |
| FC | 0.79 ± 0.05 | 0.72 ± 0.07<br>(0.75, PCA=10) | 0.90 ± 0.06 | 0.69 | 0.61<br>(0.7, PCA=15) | N = 162 |
| DKI | 0.81 ± 0.07 | 0.87 ± 0.09 | 0.79 ± 0.10 | 0.74 | 0.76 | N = 83 |
| WMTI | 0.84 ± 0.07 | 0.81 ± 0.10 | 0.82 ± 0.09 | 0.87 | 0.78 | |
| DKI+WMTI | 0.84 ± 0.06 | 0.88 ± 0.10 | 0.81 ± 0.09 | 0.82 | 0.78 | |
| FC+DKI+WMTI | 0.82 ± 0.08 | 0.77 ± 0.09 | 0.87 ± 0.09 | 0.79 | 0.76 | N = 79 |
| Ensemble<br>(FC, DKI, WMTI) | 0.85 ± 0.07 | 0.85 ± 0.09 | 0.86 ± 0.09 | 0.82 | 0.73 | |

**Figure 7**. and **Figure 8** report the SHAP value for each feature and each prediction of the LR classifiers based on DKI and WMTI parameters. As for FC, a high degree of consistency was found between metrics with high SHAP values and those displaying group differences between STZ and CTL rats. Specifically, lower FA and higher RD in corpus callosum, and lower FA and RK in fimbria were major drivers of STZ difference to CTL in the early timepoints. In the late timepoints, reduced AxD and AK in corpus callosum, decreased MK, AK and RK in cingulum, and decreased FA, MK and RK, as well as increased RD in fimbria were found to be the most prominent features distinguishing the STZ group from the CTL. WMTI-Watson parameters provided us with more specificity to differences between STZ and CTL groups. In both early and late timepoints, white mater of STZ rats was characterized by lower intra-axonal diffusivity (*D*a) in CC, indicating intra-axonal damage, and lower axonal water fraction (*f*) in CC, cingulum and fimbria, indicating demyelination and axonal loss.

**Figure 7.**
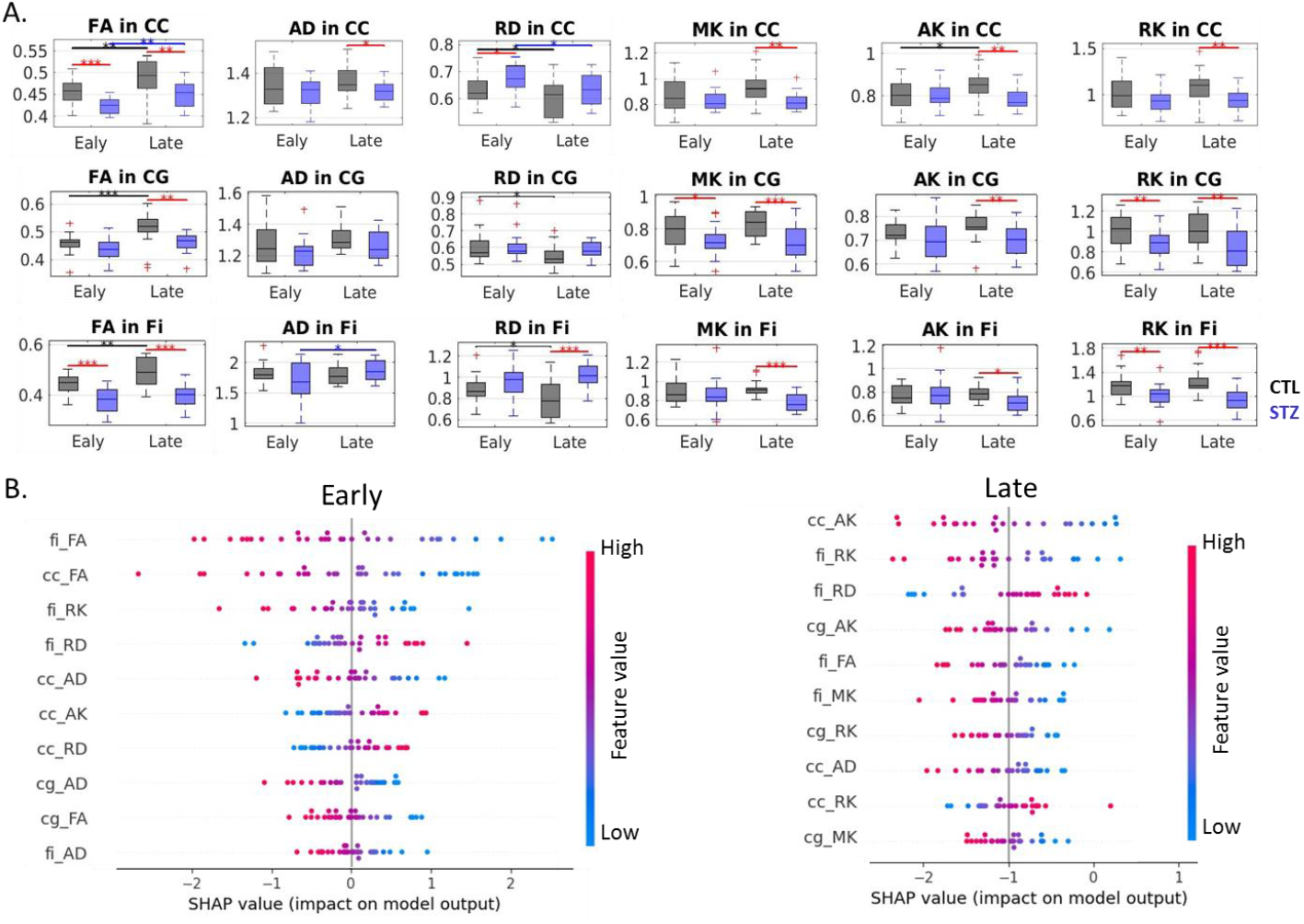
**A)** DKI estimates in three white matter ROIs (top row: corpus callosum (CC), middle row: cingulum (CG) and bottom row: fimbria of the hippocampus (Fi)). FA: fractional anisotropy, AxD/RD: axial/radial diffusivity, MK/AK/RK: mean/axial/radial kurtosis. Two-tailed *t*-test for inter-group comparison (red bars) and one-way ANOVA with Tukey-Cramer correction for within-group comparison across time (black and blue bars). ∗: *p* < 0.05, ∗∗: *p* < 0.01, ∗∗∗: *p* < 0.001. + : outlier values (but not excluded from the analysis). **B)** SHAP summary plots combining feature importance with feature effects based on DKI estimates. The position on the y-axis is determined by the feature and on the x-axis by the SHAP value. The color represents the value of the feature from low (blue) to high (red). The features are ordered according to their importance (top 10 displayed). Positive SHAP values lead the model to predict 1 (STZ) while negative ones lead the model to predict 0 (CTL).

**Figure 8.**
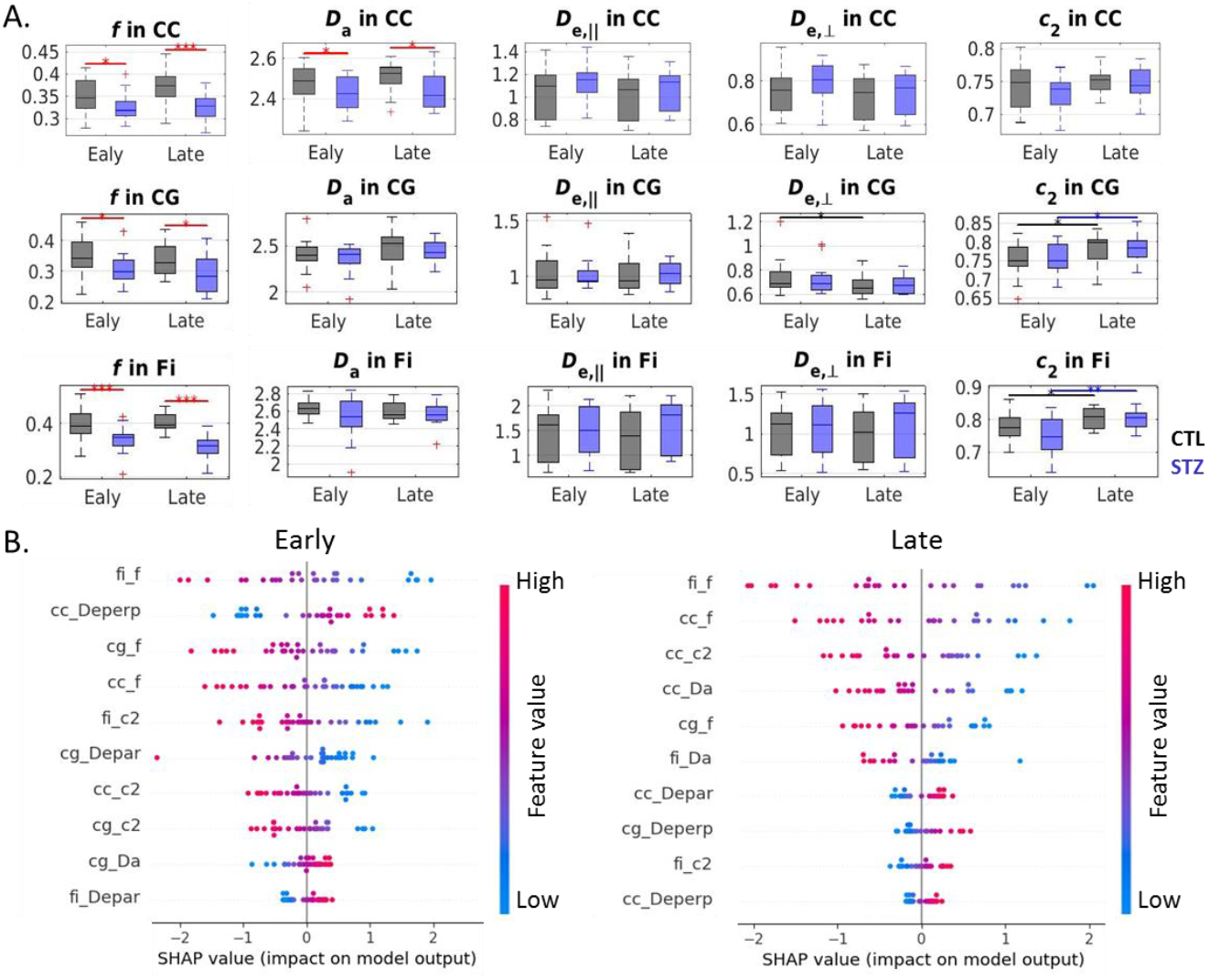
**A)** WMTI-Watson model estimates in three white matter ROIs (top row: corpus callosum (CC), middle row: cingulum (CG) and bottom row: fimbria of the hippocampus (Fi)). Two-tailed *t*-test for inter-group comparison (red bars) and one-way ANOVA with Tukey-Cramer correction for within-group comparison across time (black and blue bars). ∗ : *p* < 0.05, ∗ ∗ : *p* < 0.01, ∗ ∗ ∗ : *p* < 0.001. + : outlier values (but not excluded from the analysis). **B)** SHAP summary plots combining feature importance with feature effects based on WMTI-Watson model estimates. The position on the y-axis is determined by the feature and on the x-axis by the SHAP value. The color represents the value of the feature from low (blue) to high (red). The features are ordered according to their importance (top 10 displayed). Positive SHAP values lead the model to predict 1 (STZ) while negative ones lead the model to predict 0 (CTL).

### Combining the FC and microstructure features

After combining the FC and microstructure features, the number of total rat subjects in the pooled dataset was reduced from 83 to 79 (**Table 1**) due to the absence of either fMRI or diffusion MRI data for four datasets. The two combination methods - concatenation vs ensemble - had similar mean classification accuracy on the Late dataset, but the ensemble method achieved much higher accuracy (10% improvement) on the Early dataset, and slightly better accuracy on the Pooled dataset. Overall, neither combined classifier outperformed single classifiers at a given timepoint: best Early classification accuracy was achieved by DKI+WMTI, and best Late classification accuracy by FC.

Looking at the cross-prediction performance for all classifiers, the WMTI classifier trained on the Early dataset obtained an outstanding accuracy on the Late dataset, which was even higher than that of the classifier trained on the Late data (0.87 vs 0.82). In addition, both cross-prediction classifiers (Late-To-Early and Early-To-Late) based on WMTI features had better accuracy than those based on DKI features or combined DKI+WMTI features. The FC classifier however had poor cross prediction performance. This may indicate inter-group differences in FC evolved significantly from the early stage towards the late stage, which was consistent with early hyperconnectivity and late hypoconnectivity in STZ (**Figure 3**). The combination methods had moderate performance both in Early-to-Late and Late-to-Early predictions. With the exception of DKI, all classifiers had higher accuracy in Early-to-Late prediction than Late-to-Early.

Based on **Table 1**, a summary plot of classification accuracy based on either FC or microstructure metrics as well as the ensemble method on the three datasets (Pooled, Early and Late) is shown in **Figure 9**. On the Pooled data, the ensemble method achieved the highest overall accuracy among classifiers, which revealed that the best strategy was to combine all three types of features (FC, DKI, WMTI) in an ensemble-learning way. However, at the Early timepoint, classification based on WM microstructure, especially DKI, provided substantially higher accuracy than FC-based classification while at the Late timepoint, the FC-based classification significantly outperformed the microstructure-based classification. One possible explanation is that WM microstructure damage happens earlier than alterations in functional connectivity in the STZ group. However, this assumption needs to be further validated in human Alzheimer’s studies.

**Figure 9.**
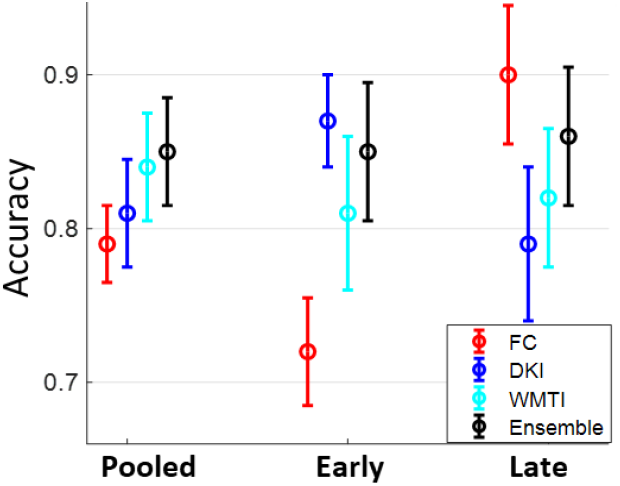
A summary plot of classification accuracy on the three datasets (Pooled, Early and Late) for each individual classifier and the ensemble classifier.

## Discussion

The classification of individuals with AD or mild cognitive impairment from healthy controls using MRI-based features and ML has been increasingly proposed. Several studies have reported promising results of employing resting-state FC as major features to this end (Hojjati et al., 2019; Ibrahim et al., 2021; Khatri and Kwon, 2022; Mousa et al., 2022; Wang et al., 2018; Wisch et al., 2020; Zhang et al., 2019). A few studies have also proposed using WM DTI-based features such FA and MD for the classification of AD subjects (Billeci et al., 2020; Doan et al., 2017; Jitsuishi and Yamaguchi, 2022; Luo et al., 2020). However, to our knowledge, this study is the first one attempting to evaluate those two types of MRI-based features seperately as well as their combination in a ML-based classification context. Furthermore, apart from the conventional DTI metrics, features based on more advanced DKI metrics and especially on biophysical models were also assessed in this study. DKI brings valuable complementary information to DTI, as seen in the list of meaningful features for classification in the Late timepoint, and the WM model narrows down the identification of microstructure changes to intra-axonal damage, demyelination and axonal loss, while providing a stable classification accuracy across time. Going forward, the acquisition of multi-shell diffusion MRI data (at least two non-zero *b*-values, e.g. *b*=1000 and 2500 s/mm^2^) in clinical studies of dementia or other brain diseases is highly recommended to enable the estimation of DKI metrics brain-wide, and of WM microstructure features using the WMTI-Watson model, for which analysis code is readily available (Diao and Jelescu, 2023).

When using FC to classify STZ/CTL rats, the most important discriminating features were connections involving the RSC, ACC, PPC, Subiculum and Hippocampus, which are regions of the default mode network typically affected by AD (Agosta et al., 2012; Brier et al., 2012; Tristão Pereira et al., 2021), as well as the hypothalamus which is responsible for recruiting alternative sources of energy to glucose, such as ketone bodies, in response to impaired brain glucose metabolism by the STZ (Carneiro et al., 2016; Foll et al., 2014; Gano et al., 2014; Le Foll, 2019; Wu et al., 2018). Many FC connections with top feature importance in the Early stage (**Figure 4**) also involved the visual and motor cortices, areas that are related to non-cognitive manifestations such as vision and motor decline and have been reported to precede the cognitive deficits (Brewer and Barton, 2014; Do et al., 2018; Hiller and Ishii, 2018; Ishii and Iadecola, 2015; Mitchell et al., 2022; Montero-Odasso et al., 2020; Vidoni et al., 2012). Moreover, the classification accuracy was improved in the Late vs Early timepoint, due to neurodegeneration progression. Finally, if only choosing connections significantly different between groups (using NBS) as features, mean classification accuracy naturally improved. In other words, connections identified as driving group differences after diagnosis could be used as features for classification in future diagnostic-blind studies.

For classification based on WM microstructure, the performance using features from a biophysical model only (here WMTI) was comparable to using all diffusion features (WMTI + DKI) and better than only using DKI features especially on the Pooled dataset. This suggests biophysical models have added value over empirical DKI tensor metrics by disengangling specific features of WM and its degeneration, and thereby classifying subjects. Microstructural features in the fimbria played the most important role in distinguishing STZ rats, which was consistent with the fact that hippocampus is especially vulnerable to AD (Mu and Gage, 2011; Setti et al., 2017) and to the icv-STZ rat model of AD. Notably, the WMTI-based classifier maintained a stable and relatively high performance across time. In fact, WMTI-Watson metrics were arguably the most stable features in discriminating STZ and CTL subjects compared to the DKI- and FC-based features, which was confirmed by the cross-prediction test where the WMTI-based classifier trained at one timepoint was more accurate in classifying subjects at a different timepoint than the other two individual classifiers. The high cross-prediction accuracy (> 0.80) might indicate the possibility of early screening and prognosis of AD in clinical applications. In other words, subjects with early WM alterations at high risk of developing further neurodegeneration might be identified and receive intervention when they are still in the early stage (van Dyck et al., 2022).

Microstructure-based features achieved better performance than FC in the Early timepoint as well as for cross-timepoint predictions. The former suggests that WM degeneration in the STZ group could happen earlier than FC breakdown. Similar findings have been reported by human studies in Subjective Cognitive Impairment as well as AD (Araque Caballero et al., 2018; Luo et al., 2020; Parker et al., 2023). The latter suggests that microstructure degeneration is relatively consistent across time. In contrast, the pattern in FC metrics was non-monotonic and shifted from Early hyperconnectivity to Late hypoconnectivity in the STZ rats, as also previously reported in human studies (Dickerson et al., 2005). However, more data are required to fully validate these hypotheses, especially in humans.

The best classification accuracy at the Early timepoint was achieved using DKI features, and at the Late timepoint using FC. However, the best overall strategy for STZ vs CTL classification was aggregating the three individual classifiers using ensemble learning. Not only was the ensemble classification more accurate on the Pooled dataset (0.85) than any of the individual classifiers, but it also maintained a high level of accuracy at each of the separate timepoints. This demonstrated that microstructural and functional information can be complementary and have their unique value in identifying STZ rats, and possibly mild cognitive impairment and early AD.

As to limitations, first, this study is based on a relatively small dataset with 24 rats followed across four timepoints. Second, we only used male rats, which was based on practical reasons. As female rats are more resistant than males to STZ-induced alterations (Biasibetti et al., 2017; Furman, 2015; Rocha et al., 2022) and hormonal modulation plays an important role in females, future studies should consider rats of both sexes. Third, in FC-based classification, each connection (ROI pair) was treated as an individual feature leading to the loss of the topological information among them. For future studies, graph neural networks can be used to replace LR for the FC-based classification (Lei et al., 2022; Zhou et al., 2020) since they consider the functional network as a whole thus better preserving spatial information. However, this will also require more advanced explainability methods to interpret the classification results (Ying et al., 2019; Zhou et al., 2022).

## Conclusions

Our work examined potential discriminators of Alzheimer’s disease in the icv-STZ rat model using functional connectivity and WM microstructure features. For the first time, we evaluated those two types of MRI-based features seperately as well as in combination, in a context of ML-based classification. WM microstructure features achieved higher classification accuracy in the early timepoints of neurodegeneration, and FC in the later timepoints, suggesting structural damage precedes functional damage. Combining all the FC and microstructure metrics in an ensemble way was the best strategy to discriminate between STZ and CTL rats, with a consistent accuracy over time above 0.85. However, for cross-time prediction, WMTI model features yielded the highest accuracy from early-to-late timepoints and vice versa, possibly thanks to the more specific metrics they capture from the microstructure, that project well across timepoints. Foreseeably in human datasets, the best microstructure (or ensemble microstructure + FC) classification features would be extracted from late timepoints with known subject diagnosis (e.g. healthy vs AD), the ML model trained on late timepoint datasets of those reduced features, and then applied to early timepoint populations to aid early diagnosis and prediction of disease evolution.

## Declarations

### Ethics approval

All experiments were approved by the cantonal and federal Services for Veterinary Affairs (VD-3306).

### Consent for publication

Not applicable.

### Availability of data and materials

The datasets generated and/or analysed during the current study are available in the OpenNeuro repository, https://openneuro.org/datasets/ds003520/versions/1.0.2 (resting-state fMRI) and https://openneuro.org/datasets/ds004441 (diffusion MRI).

## Authors’ contributions

YD collected, analyzed and interpreted the data and wrote the manuscript. IOJ designed the study, interpreted the data and edited the manuscript. All authors read and approved the final manuscript.

## Acknowledgements

The authors thank Catarina Tristão Pereira and Ting Yin for contributing code for white matter segmentation and resting-state fMRI analysis, respectively.

## Abbreviations

ACC: Anterior Cingulate Cortex
AD: Alzheimer’s disease
AxD: Axial Diffusivity
AK: Axial Kurtosis
CC: corpus callosum
CG: cingulum
CTL: control
DKI: Diffusion Kurtosis Imaging
DMN: Default Mode Network
DTI: Diffusion Tensor Imaging
FA: Fractional Anisotropy
FC: Functional Connectivity
Fi: Fimbria
fMRI: functional MRI
icv: intracerebroventricular
LR: Logistic Regression
MD: Mean Diffusivity
MK: Mean Kurtosis
ML: Machine Learning
MRI: Magnetic Resonance Imaging
PCA: Principal Component Analysis
PPC: Posterior Parietal Cortex
RD: Radial Diffusivity
RK: Radial Kurtosis
ROI: region of interest
RSC: Retrosplenial Cortex
S1: Primary Somatosensory Cortex
S2: Secondary Somatosensory Cortex
STZ: streptozotocin
WM: white matter
WMTI-Watson: White Matter Tract Integrity Watson model

